# Fidelity of and biases in the developmental acquisition of song tempo in a songbird

**DOI:** 10.1101/2022.06.17.496554

**Authors:** Logan S. James, Angela S. Wang, Jon T. Sakata

## Abstract

The temporal organization of sounds used in social contexts can provide information about signal function and evoke varying responses in listeners (receivers). For example, music is a universal and learned human behavior that is characterized by different rhythms and tempos that can evoke disparate responses in listeners. Similarly, birdsong is a ubiquitous social behavior in birds that is learned during critical periods in development and used to evoke physiological and behavioral responses in listeners. Recent investigations have begun to reveal the breadth of universal patterns in birdsong and their similarity to common patterns in speech and music, but relatively little is known about the degree to which biological predispositions and developmental experiences interact to shape the temporal patterning of birdsong. Here, we investigated how biological predispositions modulate the acquisition and production of an important temporal feature of birdsong, namely the duration of silent intervals (“gaps”) between vocal elements (“syllables”). Through analyses of semi-naturally raised and experimentally tutored zebra finches, we observed that juvenile zebra finches imitate the durations of the silent gaps in their tutor’s song with high fidelity and can “alter” the durations of gaps toward a central duration. When juveniles were experimentally tutored with stimuli containing a wide range of gap durations, we observed biases in the stereotypy but not in the central tendency of gap durations. Together, these studies demonstrate how biological predispositions and developmental experiences differently affect distinct temporal features of birdsong and highlight similarities in developmental plasticity across birdsong, speech, and music.

## INTRODUCTION

Many aspects of the temporal organization of speech and music provide important sources of information in human communication and are conserved across cultures (Ding et al. 2017; Mehr et al. 2019). For example, across languages, the duration of the silent gap (10s of milliseconds) between a consonant’s release and the onset of vocal fold vibration (known as voice-onset time) differentiates voiced consonants (e.g., /b/, /d/, and /g/ in English) from voiceless consonants (e.g., /p/, /t/, and /k/ in English; Liberman et al. 1958; Lisker & Abramson 1964; Fox et al. 2020). During conversation, interlocutors attend to and try to predict epochs of silence to coordinate the timing of their response (Zellner 1994; Stivers et al. 2009), and speakers introduce unexpected periods of silence to increase attention and arousal (Rochester 1973; Zeskind et al. 1992; Kraxenberger et al. 2018). Furthermore, variation in the temporal organization of music, including the location and duration of pauses can relate to its function and induce distinct effects in listeners (Krumhansl & Jusczyk 1990; Margulis 2007; Savage et al. 2015; Mehr et al. 2018; Bainbridge et al. 2021; Zamm et al. 2021; Mehr et al. 2021). That form-function relationships are highly conserved across cultures yet the temporal organization of speech and music is learned (e.g., Savage et al. 2015; Polak et al. 2018; Mehr et al. 2019; Roeske et al. 2020) highlights the importance of revealing the extent to which biological predisposition shape the acquisition and production of temporal patterns in speech and music.

Communication signals used by various non-human animals are structured in ways that resemble speech and music, and the temporal organization of these signals similarly can provide information (Fitch 2006; Rothenberg et al. 2014; Hoeschele et al. 2015; James & Sakata 2017; Kotz et al. 2018; Ravignani et al. 2019; Roeske et al. 2020). For example, similar to humans, epochs of silence in acoustic signaling provide information that allow interacting individuals to minimize overlap in the timing of their vocalizations (Hultsch & Todt 1982; Zelick & Narins 1983; Brumm 2006; Egnor et al. 2007; Benichov et al. 2016). On a finer temporal scale, the duration of silent gaps between distinct components of signals provides information about individual and species identity in various species, including crickets, frogs, and birds (Feng et al. 1990; Gerhardt 2005; Hedwig 2006; Coen et al. 2014; Ronacher et al. 2015; Araki et al. 2016) and can help receivers infer arousal levels and other aspects of signaler condition (Cooper & Goller 2006; James & Sakata 2014; James & Sakata 2015; Kohashi et al. 2021). Despite the similarities in the information content provided by the temporal organization of communicative elements across humans and non-human animals, relatively little is known about how biological predispositions and developmental experiences shape the temporal organization of communication signals.

Songbirds are ideal for this investigation because they learn their vocalizations during development in a manner that resembles speech and music acquisition in humans (Doupe & Kuhl 1999; Patel 2006; reviewed in Sakata & Woolley 2020). Broadly speaking, song learning in songbirds entails memorizing the sounds of a tutor’s vocalizations during a critical period in development and subsequently practicing how to reproduce those sounds. In addition, juvenile songbirds are endowed with neural processes that bias them to learn species-typical sounds and sequences (e.g., Marler & Peters 1977; Whaling et al. 1997; Gardner et al. 2005; Plamondon et al. 2008; Fehér et al. 2009; ter Haar et al. 2014; James & Sakata 2017; reviewed in Sakata & Woolley 2020). For example, when juvenile zebra finches are tutored with randomized and unbiased sequences of species-typical acoustic elements, they converge on acoustic patterns that are commonly observed in wild and laboratory populations of zebra finches (James & Sakata 2017; James et al. 2020). Songbirds like the zebra finch are also excellent to understand the temporal organization of vocalizations because activity in discrete areas of their brains regulates song tempo (Scharff & Nottebohm 1991; Long & Fee 2008; Ali et al. 2013; Zhang et al. 2017) and because sensory responses of auditory neurons are sensitive to song tempo, including the durations of silent gaps between song elements (Bee & Klump 2005; Lampen et al. 2014; Bouchard & Brainard 2016; Lim et al. 2016; Araki et al. 2016). One striking example of this temporal sensitivity is that auditory-motor neurons are most responsive to playbacks of the bird’s own song when the durations of silence are most similar to those typical of the bird (Bouchard & Brainard 2016).

To gain novel insights into the degree to which developmental experiences (learning) and biological predispositions shape the production of silent intervals within birdsong, we adopted observational and experimental approaches and quantified the precision of and predispositions in the learning of pauses in birdsong.

## METHODS

### Animals

All birds were housed on a 14 L:10 D light cycle and provided food and water *ad libitum*. All procedures were conducted in accordance to guidelines and regulations approved by the McGill University Animal Care and Use Committee in accordance with the guidelines of the Canadian Council on Animal Care. Because only male zebra finches learn and produce song, only male zebra finches were analyzed here.

#### Birds reared in a semi-natural context

The songs of 20 male zebra finches (“pupils”) and their corresponding fathers (“tutors”) from 12 nests were analyzed. Pupils raised in this manner were housed in the same nest as both parents until they were >60 days post-hatching (dph). Nests were visually occluded from each other, but songs of neighboring males were audible to pupils. Juvenile zebra finches raised in this manner (i.e., pupils can hear neighboring males’ songs but can visually, acoustically, and physically interact only with their father) generally imitate only their father’s song (Zann 1996). After 60 dph, birds were transferred to and housed in same-sex group cages and could continue to hear the songs of other zebra finches.

#### Experimentally tutored birds

Birds for experimental tutoring experiments were raised by both parents in a sound-attenuating chamber (TRA Acoustics, Ontario, Canada) up to 7 days of age, at which time their father was removed. Thereafter, juveniles were raised by only their mother. Because only male zebra finches learn to produce complex songs and because the critical period for song learning starts ^~^20 days post-hatching (Roper & Zann 2006; Brainard & Doupe 2013), this protocol ensured that juveniles were not exposed to song from a live adult during the critical period for song learning and were naïve to song before experimental tutoring. When these juveniles were nutritionally independent (i.e., could feed themselves; ^~^30-40 days old), they were housed individually in a sound-attenuating chamber for song tutoring until they were mature (^~^4 months of age; Tchernichovski et al. 2001; Derégnaucourt et al. 2005; Lipkind et al. 2013; James & Sakata 2017; Lipkind et al. 2017; Mets & Brainard 2018; Mets & Brainard 2019).

### Experimental song tutoring

Juvenile male zebra finches (“pupils”; n=67; age at start of tutoring: mean±SEM=44.6±0.8 dph; range: 31-61 dph) were individually and operantly tutored with synthesized sequences of five canonical zebra finch syllables (herein referred to as “a”, “b”, “c”, “d”, and “e”; see also James and Sakata, 2017). Each of the five syllable types was a single token taken from different birds. Zebra finch song bouts consist of a repetition of a single sequence of syllables called a “motif”, and for this experiment, pupils were tutored with a single 5-syllable motif in which each syllable appeared only once (e.g., “abcde” or “ebcda”), a common feature of zebra finch motifs (Zann 1996). We synthesized 13 motifs with different sequences and tutored, on average, 4-5 birds (range: 1-12), with each of these sequences (Table 1). Each bird was tutored with only one sequence (motif); this is because adult zebra finches sing only a single sequence of syllables that is learned during development and this form of tutoring emulates the sounds heard from a father in a nest. Zebra finch song is typically organized into bouts that contain multiple renditions of a motif, and song stimuli consisted of four renditions of a motif to emulate species-typical song bouts.

**Table 1.** Numbers of birds tutored with specific sequences (motifs) of syllables and gap durations between each syllable in the motif

|  | same gaps between syllables |  |  |  |  | different gaps between syllables | variable gaps across renditions |
| --- | --- | --- | --- | --- | --- | --- | --- |
|  | 10 | 20 | 30 | 40 | 50 |  |  |
| aecbd |  |  | n=1 |  |  |  |  |
| abdec |  |  | n=1 |  |  |  | n=3 |
| abced |  |  |  |  |  |  | n=2 |
| abdce |  |  |  |  |  |  | n=2 |
| bacde |  | n=1 | n=2 |  | n=1 |  | n=3 |
| beadc | n=1 |  | n=4 |  | n=1 |  | n=4 |
| becad |  | n=1 | n=1 | n=1 | n=1 | n=2 | n=4 |
| cdaeb | n=1 |  | n=2 |  | n=1 |  | n=4 |
| cedba |  |  |  |  |  |  | n=2 |
| dacbe | n=2 | n=1 | n=1 | n=1 | n=1 | n=1 | n=5 |
| daebc |  |  |  |  |  |  | n=1 |
| decab |  |  | n=2 |  |  |  |  |
| edbca |  |  | n=1 |  | n=1 |  | n=3 |
| eabdc |  |  |  |  |  |  | n=1 |

NOTE: The gap duration of 30 ms was overrepresented in the tutor set because previous studies of song learning used 30 ms as the set gap duration (James and Sakata, 2017); as such, tutoring birds with this gap duration provided us with the potential to directly compare learning across studies if necessary. The sequences “beadc”, “becad” and “dacbe” were also of interest based on sequencing biases (James and Sakata, 2017).

We experimentally manipulated the duration of the silent gaps between syllables in the tutor stimulus and assessed the degree to which pupils learned their gap durations from the tutor stimulus. One group of birds (n=30) was tutored with a tutor sequence in which gap durations were fixed to be either 10, 20, 30, 40 or 50 ms (“stereotyped gap stimulus”; 10 ms: n=4; 20 ms: n=3; 30 ms: n=15; 40 ms: n=2; or 50 ms: n=6; Table 1). These durations were chosen based on observed distributions of gap durations within our (and other lab’s) colony of birds (e.g., Price 1979; Norton & Scharff 2016; James & Sakata 2017); for example, the vast majority of gaps within motifs are <50 ms, with peaks around 30-40 ms (Cooper & Goller 2006; Norton & Scharff 2016; Lachlan et al. 2016; Araki et al. 2016; see also Fig 1). Additionally, this range of gap durations was selected based on the gap duration (30 ms) used in a previous study investigating biases in a feature of the temporal organization of song, namely syllable sequencing (James & Sakata 2017). The differences in gap durations are likely to be conspicuous to the birds since songbirds (including zebra finches) are sensitive to differences of 1-2 ms of sounds (Dooling et al. 2002). In each of the stereotyped gap stimuli, the silent interval was the same between each syllable in the motif (“within-motif gap”) and across each rendition of the motif. For example, if a bird was tutored with the sequence “abcde” with a 30-ms gap, the gaps between “ab”, “bc”, “cd”, and “de” were 30 ms for each rendition of the sequence. The gap between syllables across adjacent motifs (i.e., from offset of last syllable in the motif to onset of first syllable of subsequent motif: “between-motif gap”) was kept constant at 100 ms across bouts and birds (see James and Sakata, 2017). Longer gaps between motifs compared to gaps within motifs is characteristic of zebra finch song (Vu et al. 1994; Glaze & Troyer 2006; Hyland Bruno & Tchernichovski 2019).

**Figure 1:**
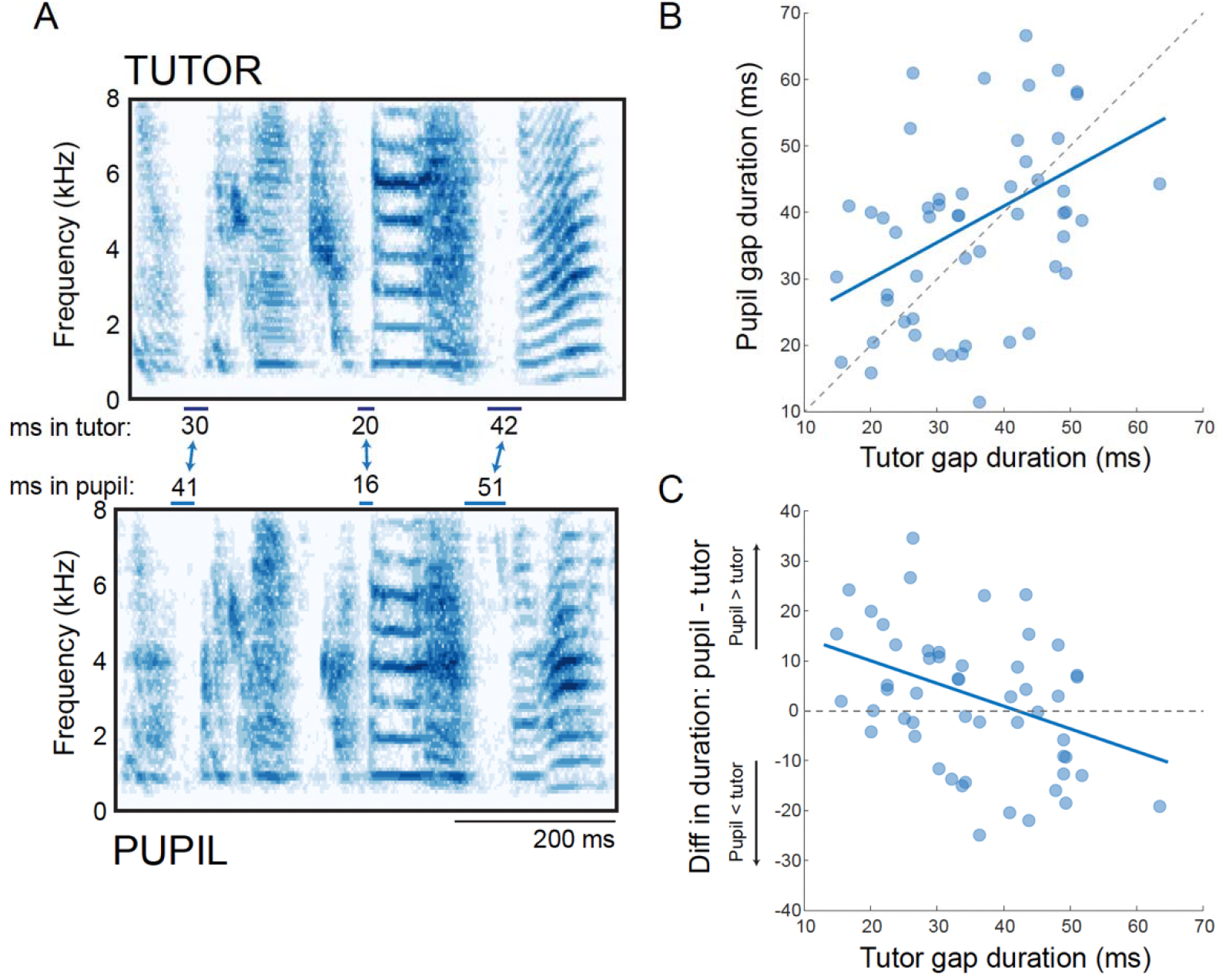
Learning of gap durations in a naturalistic context. A) Example of gap learning between a tutor and his pupil. The tutor’s motif is on top and below is an excerpt of the pupil song that contains all three of the learned gaps from the tutor. [One syllable in the tutor’s song (the 2^nd^ syllable) was “split” by the pupil, creating a novel gap that was not considered to be shared with the tutor (see also Tchernichovski et al. 2021).] B) Positive correlation between tutor and pupil gap durations (n=52 shared gaps). Lines indicate the line of unity (dashed) and the significant trendline from a mixed-effects model (solid). C) Negative relationship between the gap duration in the tutor and the deviation of the pupil’s gap duration from the tutor’s gap duration (i.e., difference in duration between the tutor and pupil). Lines indicate y = O (dashed) and the significant trendline from a mixed-effects model (solid).

Three birds were also tutored with gap durations that did not vary across renditions of the motif but in which gap durations varied between different syllables in the motif. For example, two birds were tutored with the motif sequence “becad” and one bird with the sequence “dacbe” but in which the four gaps between syllables within the motif were fixed, respectively, at 20, 50, 10, and 40 (Table 1). These birds are integrated into the analysis of birds tutored with stereotyped gap durations (for both sets of birds, the gaps were stereotyped across renditions of the motif; n=33 birds in total).

Another group of birds was tutored with a stereotyped sequence of syllables (i.e., a single motif) but in which the durations of each silent gap between syllables within the motif (within-motif gap) randomly varied between 10-50 ms across renditions of the motif (n=34 birds; in discrete 10-ms steps: 10, 20, 30, 40, or 50 ms; “variable gap stimulus”). For example, a bird could be tutored with the sequence “abcde” and on some renditions of the sequence the gap between “a” and “b” was 10 ms, and on other renditions it was 20, 30, 40, or 50 ms. Stimuli were synthesized such that the gap duration was pseudorandomly determined and that birds were exposed to each of the five gap durations at equal probabilities. Tutoring birds with such ambiguous stimuli enables the discovery of biases in vocal learning (e.g., James and Sakata, 2017). This degree of variability in gap durations within the motif was substantially higher than that observed in natural zebra finch song; the normalized IQR (IQR/median) of gaps within the variable gap stimulus was 0.67, compared to 0.12 for the average normalized IQR of gaps within motifs of semi-naturally reared birds in this study. However, similar to stimuli with stereotyped gap durations within the motif, the between-motif gap was kept constant at 100 ms across bouts and birds. Twelve different sequences were used for variable gap stimuli (Table 1).

Tutored birds were operantly tutored using perch hops for >1 month (mean±SEM=47.4±1.3 days); this method and duration of tutoring is comparable to previous studies (ten Cate 1991; Adret 1993; Derégnaucourt et al. 2013; James & Sakata 2017; Mets & Brainard 2018; Mets & Brainard 2019). Song playback was operantly triggered by perch hops (n=65) or string pulls (n=2) using custom-built perches or strings connected to a National Instruments PCI-6503 I/O card (National Instruments, TX), with each hop or string pull triggering the playback of one song bout (i.e., four concatenated motifs of the tutor stimulus). Song playbacks were spaced in time such that juveniles could hear only 10 operantly triggered song bout playbacks within each of three time periods in the day (morning, noon, and afternoon); this schedule of song exposure has been found to enhance song learning (Tchernichovski et al. 1999; Derégnaucourt et al. 2005). There was no significant difference between birds tutored with stereotyped or variable gap durations in the age at which tutoring began (stereotyped gaps: 45.3±1.2; variable gaps: 44.0±1.0; t-test: t_65_=0.8; p=0.4161) or in the duration of tutoring (stereotyped gaps: 48.5±2.0; variable gaps: 47.6±1.6; t-test: t_65_=0.4; p=0.7136). Sound Analysis Pro 2011 (SAP; http://soundanalysispro.com) was used for song tutoring, and stimuli were played out of an Avantone Pro Mixcube speaker (Avantone, NY) connected to a Crown XLS 1000 amplifier (Crown Audio, IN).

### Song recording

For all song recordings, birds were housed individually in a sound-attenuating chamber (TRA Acoustics, Ontario, Canada), and song was recorded using an omnidirectional microphone (Countryman Associates, Inc, Menlo Park, CA) positioned above the bird’s cage. Computerized, song-activated recording systems were used to detect and digitize song (SAP; digitized at 44.1 kHz). Recorded songs were digitally filtered (0.3–10 kHz) for off-line analysis using software custom-written in the Matlab programming language (MathWorks, Natick, MA). All songs recorded and analyzed were spontaneous songs produced in isolation (“undirected song”).

For our analyses of song learning, all pupils were recorded as adults (>90 dph). For the dataset of colony-reared birds, we analyzed the recordings of adult tutors that were as close as possible to the recording date of pupils.

### Identifying song motifs and shared gaps

We analyzed the adult songs of colony-reared or experimentally tutored zebra finches. We identified and labeled syllables following amplitude-based segmentation of audio files using custom-written MATLAB scripts [14.3±0.6 (mean±SEM) songs per bird]. The song of an adult zebra finch generally consists of a single stereotyped sequence of syllables called a “motif”. An individual’s motif is readily identifiable because it is repeated multiple times within a song bout (Scharff & Nottebohm 1991; ten Cate 1991; Zann 1996; Tchernichovski et al. 2000) and because the duration of gaps between motifs is usually longer than gap durations within the motif (Vu et al. 1994; Glaze & Troyer 2006). To identify motifs in the songs of tutors (n=12), three authors (LSJ, ASW & JTS) independently examined multiple renditions of each tutor’s song and identified motifs based on sequence repetition, sequence stereotypy and gap durations. Because only gaps within the motifs of experimental stimuli were manipulated (see below), we only analyzed the gaps that were produced within the motif of the tutor or tutoring stimulus.

The main sets of analyses covary gap durations in the tutor or tutoring stimulus (i.e., the tutor’s song or tutoring stimulus) with the duration of the same gap (“shared gap”) in the pupil. We asked a rater (J. Thompson) who was familiar with zebra finch song but naïve to the experiment to identify shared gaps between pairs of spectrograms. For this, the rater viewed an image with spectrograms of two songs and was asked to identify gaps that were shared across songs by highlighting epochs of silences that occurred between acoustically similar sounds. The rater viewed all tutor-pupil or tutoring stimulus-pupil pairs at least twice, and we included each gap that was identified as shared in our analyses.

We computed a variety of metrics to assess the validity of the approach (Fig S1). (1) We asked the rater to identify shared gaps between 15 pairs of syllable sequences that were synthesized using the same syllable tokens for experimental tutoring (positive controls); in this dataset, we know the number of syllable pairs (sequences) that are shared or distinct between pairs of synthesized songs. Regardless of the gap durations in each of the songs that were compared, the rater identified all of the shared gaps (22 out of 22) and only the shared gaps (i.e., did not misclassify any gaps as “shared”). For example, the sequences “abdec” and “abdce” share two common gaps (“ab” and “bd”) that were both identified by the rater; on the other hand, the sequences “beadc” and “decab” share no common gaps and none were identified as shared by the rater. (2) The rater was also presented with “distractor” pairings in which a spectrogram of a song by a tutor in our breeding colony was randomly paired with a spectrogram of a non-pupil (n=20 pairings). Shared gaps were noted in only 1 of the 20 random pairings, which can be expected given some features of song are shared across birds in the colony. (3) We computed the intra-observer reliability in the dataset of colony-reared birds by providing the rater with replicate pairings and computed the proportion of gaps that were consistently identified as shared across the duplicate pairings. We presented two types of replicate pairings: (a) the exact same song exemplars for two birds were presented on two different images and (b) two different song examples from the same two birds were presented on different images. Combining across these two types, 89% (58 out of 65) of gaps were consistently identified as shared across replicates.

### Statistical analyses

After the identification of shared gaps, we computed the median and normalized IQR (“normIQR” = interquartile range divided by the median; analogous to the coefficient of variation) as the measures of the central tendencies and variation of each gap. To compare the durations and variability of gaps that were shared between tutors and their pupils in our semi-natural breeding context, we used a mixed-effects model with the pupil gap duration as the dependent variable and the tutor gap duration as the independent variable. Because each tutor can have multiple pupils and because each pupil can contribute multiple data points (i.e., multiple gaps within a pupil’s song can be shared with the tutor), we included both tutor ID and pupil ID as random effects in the model. The same model was used to analyze similarities in the variability of gap durations between tutors and pupils. To analyze gap durations in pupils experimentally tutored with stereotyped gap durations, we used a similar model with the gap duration in the tutor stimulus as the independent variable, and pupil ID as a random effect. Similar mixed effects models were used to compare the normIQRs across methods of tutoring.

Finally, for birds experimentally tutored with variable gap durations, we used *χ*^2^ tests as well as permutation tests (Monte Carlo simulations) to assess whether the observed distribution of gaps differed from that expected by chance. Gap durations were converted into bins of equal durations centered at the durations used for tutoring (10, 20, 30, 40 and 50 ms); specifically, the bin intervals for this analysis were: 5.0-14.9, 15.0-24.9, 25.0-34.9, 35.0-44.9, and 45.0-54.9 ms. Observed data were binned in this manner and converted into proportions. The expected distribution was an even distribution of gap durations across five bins (i.e., null hypothesis that birds are equally likely to produce any one of the five gaps in the tutor stimulus).

For the permutation, we randomly selected (with replacement) one of the five tutor gaps and created a distribution of permuted gaps (of the same sample size as the dataset). The distribution of permuted gaps was converted into proportions and used to compute various metrics for analysis. This was repeated 1000 times to create a distribution of permuted metrics.

For the permutation test, we computed two metrics. First, we analyzed how the spread of the distribution of observed gaps compared to the spread of the distribution of expected gaps. To this end, we first computed the entropy of the observed distribution of proportions (“observed entropy”) and the entropy of the expected distribution of proportions (“expected entropy”), and then calculated the difference between the observed and expected entropy (“observed entropy difference”; observed entropy minus expected entropy). Because the expected distribution is an even distribution, the expected entropy is the maximum entropy; as such, the observed entropy difference was negative. For each iteration of the permutation, we computed the entropy of the permuted distribution (“permuted entropy”) as well as the difference between the permuted and expected entropies (“permuted entropy differences”). Finally, we asked how often one would observe a difference as negative as the observed entropy difference in the dataset of permuted entropy differences.

Second, we computed the Jensen-Shannon divergence (JSD) between the observed and expected probability distributions. The JSD is a metric that quantifies the difference between two probability distribution and is related to the K-L divergence. To assess the significance of the observed JSD, we computed the JSD between the permuted and expected distributions on each iteration of the permutation and then quantified how likely it was to observe, by chance, a JSD as large as the observed JSD.

## RESULTS

### Gap learning in a naturalistic context

We first analyzed the extent of similarity in gap durations between 20 tutor-pupil pairs (from 12 nests) in a colony of semi-naturally breeding zebra finches. The average duration of shared gaps within the motifs of tutors and pupils in our colony was 36.0±1.4 (mean±SEM) ms (*n* = 89 gaps; combined data from pupils and tutors), which is comparable to values reported in other studies (e.g., Glaze and Troyer, 2006; Lachlan et al., 2016; Araki et al., 2016).

We identified 52 gaps within the motifs of tutors that were reproduced by their pupils (“shared gaps”; see Methods). An example of the songs of a tutor and his pupil are provided in Figure 1A, and shared gaps are highlighted. In this example, the pupil learned all three of the tutor’s gaps, and reproduced them with similar gap durations. Across all tutor-pupil pairs, the durations of shared gaps were not significantly different between tutors and pupils (*F*_1,51_ = 0.9, *p* = 0.3381). Moreover, gap durations in the motifs of tutors were significantly correlated with the corresponding (shared) gaps in pupils (*F*_1,41.4_ = 11.5, *p* = 0.0016; Fig 1B).

We also observed an interesting pattern for gap durations, where pupils tended to lengthen relatively short gaps in their tutor’s song and shorten relatively long gaps in their tutor’s song (i.e., the relationship was not 1:1). Correspondingly, there was a negative relationship between the gap duration in the tutor’s motif and the deviation of the pupil’s gap from the tutor’s gap (pupil duration minus tutor duration; *F*_1,41.4_= 8.0, *p* = 0.0072; Fig 1D). Specifically, gaps <30 ms in duration in the tutor’s song tended to be longer in the pupil’s song compared to the tutor’s song, and gaps ≥40 ms in the tutor’s song tended to be shorter in the pupil’s song compared to the tutor’s song. This pattern of differences between the tutor and pupil’s song is consistent with a regularization process and suggests that biological predispositions could shape the learning of gap durations.

### Gap learning in an experimental context

To control for genetic, social and other environmental factors that may have influenced gap learning in a naturalistic context (wherein tutors are the fathers of pupils), we experimentally tutored song-naïve zebra finches (i.e., raised without song exposure during the critical period of song learning; see Methods). Each pupil was tutored with a synthesized stimulus consisting of a stereotyped sequence of syllables (motif) in which the duration of gaps between syllables in the motif was fixed between 10-50 ms (in 10-ms increments; see Methods; Table 1; “stereotyped gap durations”), a range of durations typically found in zebra finches (see also Fig 1B). In other words, gap durations between pairwise sequences of syllables were fixed to the same duration across each rendition of the motif (Fig 2A).

**Figure 2:**
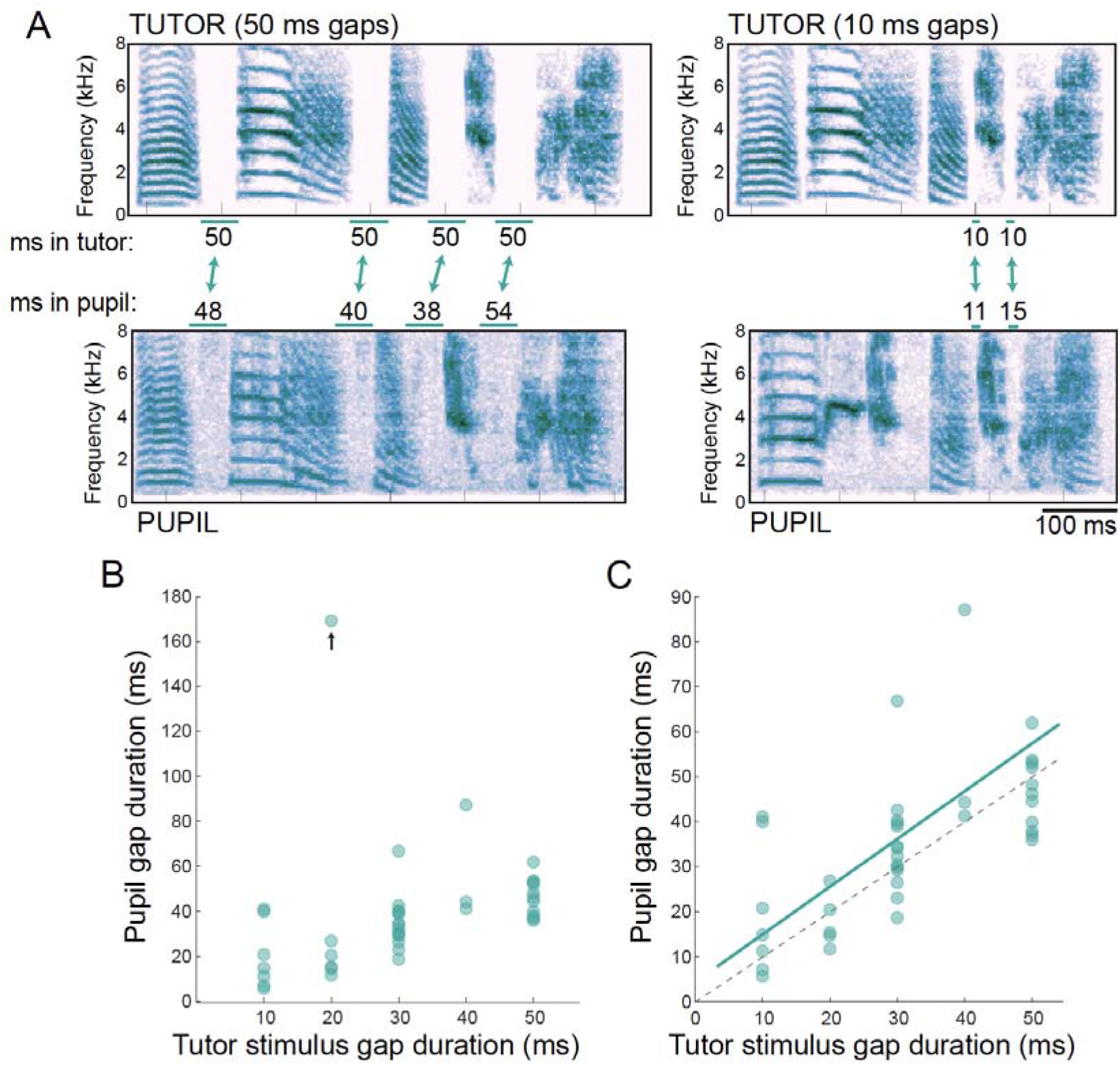
Learning of gap durations in tutor stimuli with stereotyped gaps. A) Examples of gap learning between pupils (bottom) and their tutor stimuli (top). These two pupils were brothers and were tutored with the same tutor sequence, but different gap durations (10 vs. 50 ms gaps). B) Correlation between the gap in the tutor stimulus and the shared gap in the pupil. Arrow indicates outlier. C) Same correlation as (B), plotted without the outlier. Lines indicate the line of unity (dashed) and the significant trendline from a mixed-effects model (solid).

We identified 41 gaps in the songs of 24 pupils that were shared with gaps within the motifs of the tutor stimulus. Two examples of pupils that were tutored with the same sequence of syllables but with different gaps between syllables are provided in Figure 2A; in one example, the gaps between all syllables in the motif were fixed to 50 ms (left stimulus), whereas in the other example, the gaps between the syllables in the motif were fixed to 10 ms (right stimulus). Gaps that were identified to be shared between the pupil’s song and the tutor stimulus were longer for the bird tutored with 50 ms gaps than for the bird tutored with 10 ms gaps. Across all pupils, we observed a marginally significant correlation between the durations of shared gaps in the tutor stimulus and the pupil’s song (*F*_1,39_ = 4.0, *p* = 0.0532; Fig 2B). However, the lack of statistical significance was driven by one data point, and this relationship was highly significant following the removal of the outlying point (*F*_1,27.6_ = 59.8, *p* < 0.0001; Fig 2C). Moreover, the slope of this relationship was nearly identical to 1 (estimate±SEM: 1.06±0.14), and the intercept was not significantly different from 0 (*p* = 0.4020).

### Biases in gap learning and production

To experimentally reveal the degree to which biological predispositions sculpt the acquisition and production of gap durations, we individually tutored song-naïve zebra finches with randomly varying gap durations. Specifically, each pupil was tutored with a stereotyped sequence of syllables (motif) in which the duration of gaps between syllable pairs within the motif randomly varied between 10-50 ms (in 10-ms increments) across every rendition (see Methods). We hypothesized that if biological predispositions to produce species-typical gap durations (e.g., 30-40 ms) exist, we should observe a distribution of gap durations that was significantly different from a uniform distribution (null distribution).

A rendition of an adult pupil’s song and a rendition of his tutor stimulus are depicted in Figure 3A. Two motifs in the tutor stimulus are depicted, and the duration of the gaps between syllables (e.g., between the first and second syllables of the motif) differ between the two motifs (highlighting the variability of gap durations in the stimulus). In this example, all four gaps in the stimulus motif were identified as shared in the pupil’s song. Despite being tutored with variable gap durations, the adult song of the pupil consisted of gaps that were relatively consistent in duration across renditions (see consistency of gaps between motifs in the pupil’s song; see below for analyses of variability), and for this bird, the durations of the shared gaps were 11, 40, 36, and 40 ms. Across all birds tutored in this manner, we identified 50 total shared gaps across 24 birds. The average duration of shared gaps was 32.8±2.5 (mean±SEM) ms (Fig 3B). To analyze the distribution of gap durations in these tutored birds, we binned the data based on the tutor stimulus (e.g., 10 ms bins with the first 5 centered at 10, 20, 30, 40 and 50 ms). We found no evidence that a particular gap duration was favored (Likelihood ratio test: *χ*^2^_4_=3.9, p=0.4131; Fig 3C), suggesting minimal or weak biases to produce specific gap durations. However, gap durations ≥45 ms were relatively rare (Fig 3C), suggesting a general trend for gaps to be less than 45 ms.

**Figure 3:**
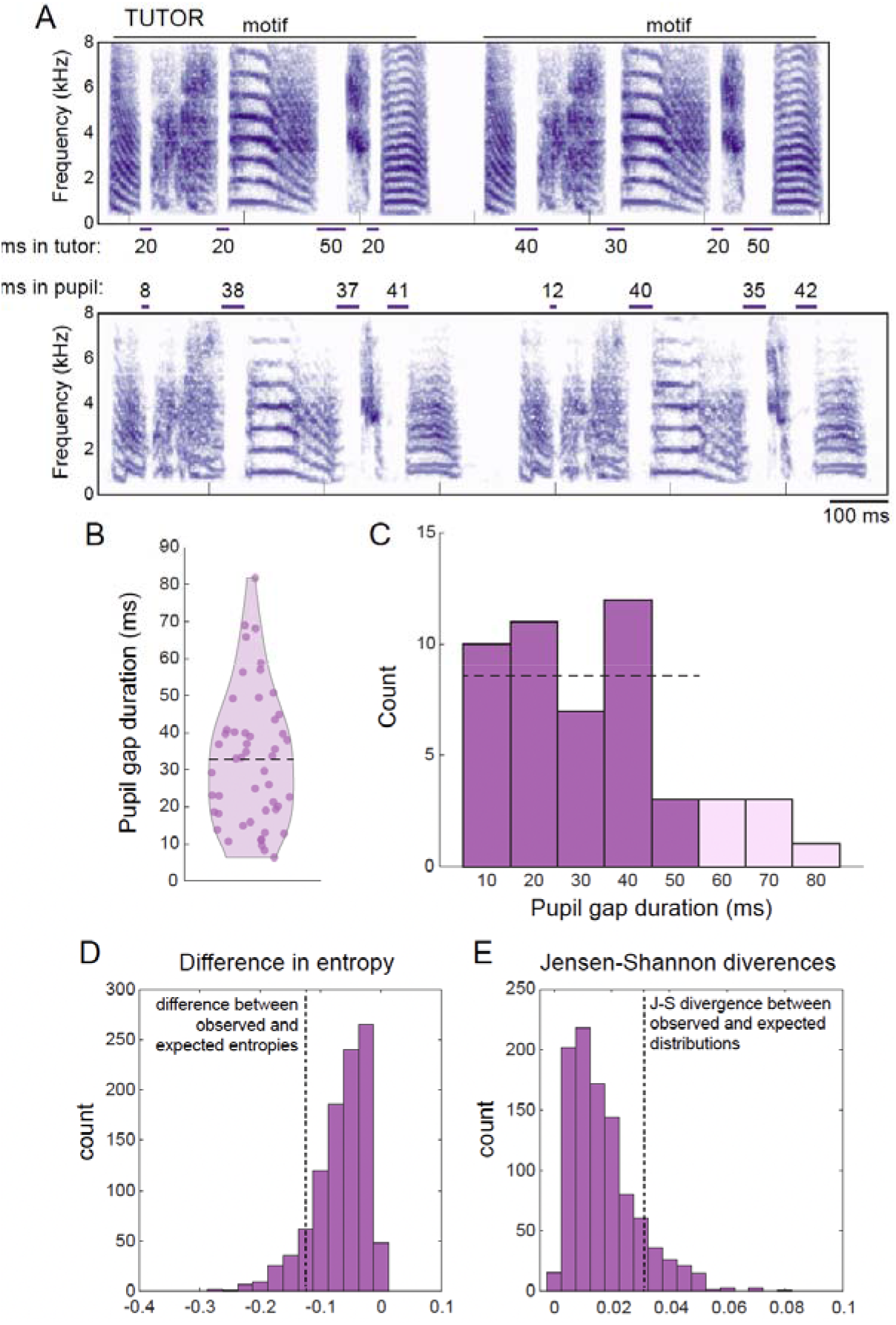
Learning of gap durations in tutor stimuli with variable gaps. A) Example of a tutor stimulus (top) and the pupil’s song (bottom). Note that, from motif to motif, the durations of gaps are variable in the stimulus, but relatively stereotyped in the pupil. This pupil reproduced all four gaps within the motif of his tutor stimulus. B) Violin plot for all gaps produced by pupils that are shared with the tutor stimulus. Each dot represents one gap, and the dashed line indicates the mean. C) Same data as (B), in 10 ms bins. The first five bins (dark bars) are centered on each of the five gap durations present in the tutor stimulus. The dashed line indicates the null distribution expected based on the tutoring stimuli (equal number of gaps in each bin). D-E) Histograms of the outputs from permutation tests measuring the entropy of the distribution of gaps (D) and the Jensen-Shannon divergence (E). Vertical dashed lines indicate the observed values.

We used permutation tests (Monte Carlo simulations; see Methods) to further assess the significance of the distribution of gap durations produced by birds tutored with randomized gaps. We first analyzed the entropy (spread; measured in bits) of the distribution of observed gap durations and compared this to the entropy of the distribution of expected gap durations (the expected distribution is an even distribution of gap durations across the five bins of gap durations; Fig 3D). The observed entropy difference was −0.1213 bits (observed minus expected). We then asked how likely it was to observe a difference as negative as the observed difference in the permuted dataset, and found that 11.7% of permuted differences were equal to or less than the observed difference (i.e., p=0.1170). We also computed the Jensen-Shannon divergence (JSD) between the observed and expected probability distributions (observed JSD: 0.0331) and found that JSDs equal to or greater than the observed JSD were found on 10.2% of the permutations (p=0.1020; Fig 3E). Consequently, these randomization tests also support the notion of minimal or weak biases in the learning of gap durations.

### The relationship between gap and syllable durations

Experimentally tutoring pupils with randomized gap durations allows the investigation of other song features that may influence gap durations. For example, because zebra finches generally inhale during the gaps between syllables (“mini-breaths”; Hartley & Suthers 1989; Franz & Goller 2002; Cooper & Goller 2006), longer gaps may be associated with longer syllables before or after the gap (e.g., Mota & Cardoso 2001; Zollinger & Suthers 2004; Cardoso et al. 2007). However, we observed minimal evidence for a relationship between syllable and gap durations for birds tutored with random gap durations (durations of the gap and the syllable <u>before</u> the gap: *F*_1,24.5_ = 3.1, *p* = 0.0914; Fig 4A; durations of the gap and the syllable <u>after</u> the gap: *F*_1,48_ = 0.2, *p* = 0.6900; Fig 4B).

**Figure 4:**
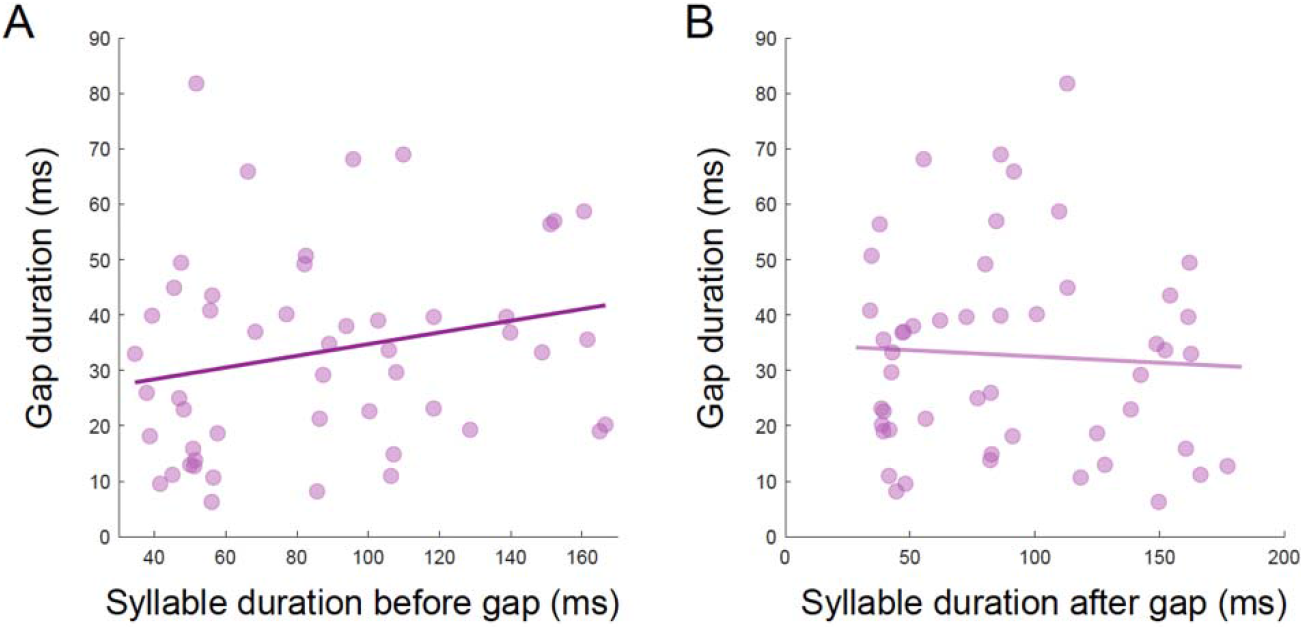
For birds tutored with variable gap durations, the durations of gaps were not significantly related to the duration of the syllable before (A) or after (B) the gap.

### The relationship between the variability of gap durations in the tutor and pupil’s songs

The variability of some temporal features of song (e.g., the variability of syllable sequencing) is learned (e.g., James et al., 2020), but the degree to which variability in gap durations (syllable timing) is learned remains unknown. In our analysis of semi-naturally reared birds, the variability of gap durations (normIQR) was not significantly correlated between pupils and tutors (*F*_1,45.2_= 1.6, *p* = 0.2070). When comparing the variability of shared gap durations in birds experimentally tutored with stereotyped vs. variable gap durations, shared gap durations were equally variable between these two sets of experimentally tutored birds (*F*_1,36.2_ = 0.1, *p* = 0.7910). Both groups of experimentally tutored birds produced gaps whose variabilities were not significantly different from those birds tutored under semi-natural conditions (*F*_2,59.0_ = 2.5, *p* = 0.0888). These results are particularly striking for birds tutored with variable gap durations, where the input (tutor stimulus) contained atypical high variability in gap durations but the birds produced gaps with species-typical variability.

## DISCUSSION

Just as intervals separate notes within music or phrases within speech, intervals of silence (“gaps”) separate acoustic units within the songs of songbirds. Here, we manipulated the developmental experiences of juvenile zebra finches to reveal the degree to which learning and biological predispositions shape the acquisition of gap durations in the zebra finch. Birds that were reared in a semi-natural setting (and tutored by their father) generally learned the gap durations of their tutor (father), but also tended regularize the longest and shortest gaps in their tutor’s song towards the mean. Birds experimentally tutored with stereotyped gap durations also imitated the gap durations present in the stimulus, confirming the importance of learning for gap durations. Finally, birds experimentally tutored with stimuli containing variable gap durations generally produced songs with a range of possible gap durations, highlighting the flexibility in the acquisition and production of gaps.

Broadly speaking, the observation of significant developmental learning of gap durations in zebra finches is consistent with previous studies documenting the neural control of gap durations by brain areas important for vocal learning. For example, reinforcement paradigms can drive changes to the duration of gaps between syllables in adult finches, and this plasticity is influenced by activity within a cortical-basal ganglia loop that is critical for developmental song learning (Ali et al. 2013; Tachibana et al. 2017). In addition, the current data from semi-naturally raised zebra finches are consistent with the acquisition of gap durations in cross-fostered zebra finches. In particular, Araki et al. (2016) cross-fostered juvenile zebra finches with adult Bengalese finches and observed a positive relationship between gap durations of adult Bengalese finch tutors and their zebra finch pupils. In addition, they found that, while gap durations <100 ms were accurately imitated by zebra finch pupils, gaps >100 ms tended to be truncated by juveniles. We observed similar truncation of relatively long gaps in a conspecific tutor’s song among semi-naturally raised zebra finches (Fig 1), suggesting that the regularization is not simply due to cross-fostering. Combining the results across experiments, these data suggest that it will be important to reveal the degree to which biases and accuracies in developmental gap learning are shaped by activity within the cortical-basal ganglia loop for vocal plasticity or in auditory circuits (e.g., Ali et al. 2013; Araki et al. 2016).

Interestingly, experimental tutoring did not provide strong support for biological predispositions in the acquisition and production of gap durations. For example, neither the truncation of tutor gaps >40 ms nor elongation of tutor gaps <30 ms was observed in zebra finches that were tutored with stereotyped gap durations (Fig 2). The observation of gap “regularization” in semi-naturally and socially-reared birds but not in birds that were individually tutored with song playbacks seems counter to previous studies of learning. In particular, because social interactions promote the learning of syllable structure (Chen et al. 2016; Ljubičić et al. 2016; reviewed in Sakata & Yazaki-Sugiyama 2020), one might assume greater fidelity of gap learning (i.e., similarity between pupil and tutor’s songs) in socially tutored birds than in operantly-tutored birds. However, median gap durations of socially tutored birds differed in a more systematic manner from their father’s (tutor’s) song than birds that were experimentally (operantly) tutored (Figs 1 and 2). A significant truncation of relatively long gaps could be observed if longer gaps (>50 ms) were used in the tutor stimulus; nevertheless, the elongation of relatively short gaps was not observed in birds experimentally tutored with stereotyped gaps. It is possible that the greater stereotypy of gap durations across renditions and increased consistency of gaps within motifs for the tutor stimulus and/or variation in the amount of song exposure during development contributed to this difference between semi-naturally and experimentally tutored birds.

Minimal evidence of biases in gap learning was observed in zebra finches that were experimentally tutored with a randomized and unbiased distribution of gap durations. Given the prevalence of gap durations ^~^30-40 ms in published studies (e.g., Scharff & Nottebohm 1991; Norton & Scharff 2016; Araki et al. 2016) and regularization observed in semi-naturally reared birds, we anticipated that birds tutored with a wide range of gap durations would significantly “prefer” to produce gaps between tutor syllables that were 30-40 ms in duration. However, many shared gaps were <30 ms in duration, and while there was a trend against producing relatively long gaps, the overall distribution of observed gap durations was not statistically different from a uniform distribution (i.e., the distribution of gaps birds were tutored with). Even though the ranges of gaps used for experimental tutoring were comparable to those observed in the social tutors of semi-naturally reared birds (Fig 1), it remains possible that tutoring birds with a wider range of gap durations might more effectively demonstrate biases in gap learning.

The lack of strong biases in the acquisition of gap durations contrasts with the significant biases in the acquisition of another temporal feature of song, namely syllable sequencing. Just as studies of zebra finch song demonstrate the prevalence of gaps that were 30-40 ms in duration, studies of wild or laboratory populations of zebra finches observed common sequences of syllables (Zann 1996; Lachlan et al. 2016). For example, short, high-pitched syllables are often produced in the middle of the motif sequence whereas longer syllables resembling a “distance call” are often placed at the end of the motif sequence. These observations suggested the possibility of biases in syllable sequence learning. James and Sakata (2017) individually tutored juvenile zebra finches with randomized and unbiased sequences of five common, species-typical syllables (same syllables used in the current study) and observed convergent patterns that matched those commonly found in wild populations of zebra finches. In addition, the acoustic patterns produced by birds tutored with randomized sequences resembled patterns frequently observed in music (e.g., phrase-final elongation, harmonic arches, and pitch alternation). Collectively, these data suggest biases in sequence learning could be stronger than biases in acquisition of syllable timing and suggest distinct mechanisms underlying the acquisition and control of these temporal features of song. Indeed, some previous studies have discussed both shared and distinct mechanisms underlying syllable sequencing and timing (Hampton et al. 2009; Glaze & Troyer 2013; Matheson & Sakata 2015; Troyer et al. 2017).

A number of these findings about gap learning in zebra finches have parallels with investigations of music acquisition and imitation. For example, just as people can accurately reproduce musical notes at various intervals and tempos, juvenile zebra finches can accurately reproduce gap durations heard in during development. In addition, the relatively weak bias for gap learning compared to sequence learning in zebra finches suggests a dissociation between the acquisition of sequencing and timing in songbirds, just as reproductions of the sequence and timing of notes or movements are also partially dissociable in humans (e.g., Pfordresher 2003; Ullén & Bengtsson 2003; Brown et al. 2013; Kornysheva et al. 2013; Kornysheva & Diedrichsen 2014). Moreover, the propensity for zebra finches tutored with random gap intervals between syllables to produce songs with relatively stereotyped gap durations is similar to findings of beat generation in humans. Music is typically produced at regular intervals (rhythms), and when humans are tasked to replicate a stimulus with irregular intervals between beats, participants tend to “regularize” the beats into predictable patterns over-time (Ravignani et al. 2016; Jacoby & McDermott 2017). Given these parallels between the temporal organization and plasticity of music and birdsong, these data suggest that songbirds offer a powerful opportunity to test hypotheses about the neural substrates underlying the development and organization of music, including rhythm.

## Data availability statement

The data that supports the findings of this study will be made available in the supplementary material of this article.

## Conflict of interest statement

The authors have no conflict of interest to declare.

## Funding statement

This work was supported by funding from the National Science and Engineering Research Council (#05016 to J.T.S.), the Fonds de Recherche du Quebec - Nature et technologies (PR-299652 to S.C.W.), the Centre for Research on Brain, Language and Music (L.S.J.), and a Heller award (L.S.J.).

## Ethics approval statement

All procedures were conducted in accordance to guidelines and regulations approved by the McGill University Animal Care and Use Committee in accordance with the guidelines of the Canadian Council on Animal Care.

## Acknowledgements

We would like to thank J. Thompson for help with song annotation, M. Bertolo for discussions about song annotation and analysis, M.H. Kao for discussions about data, and S.C. Woolley for comments on manuscripts and data analysis.

---

**Figure S1.**
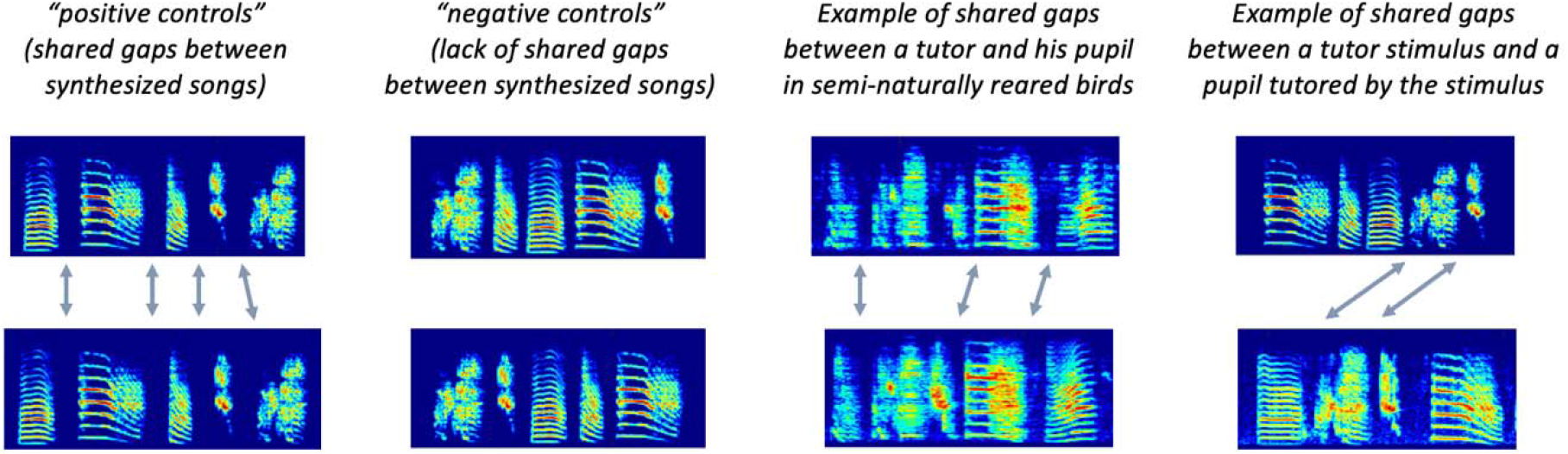
Examples of shared gaps identified by a rater (J. Thompson). Depicted are, from left to right, an example of shared gaps identified in a positive control (pairs of synthesized stimuli with known shared gaps), of the lack of gaps identified as shared in a negative control, of shared gaps between a tutor and his pupil in semi-naturally reared birds, and of shared gaps between a tutor stimulus and a pupil that was experimentally tutored with the stimulus. All pupil songs were recorded when they were adults (see Methods).

## Notes

### Competing Interest Statement

The authors have declared no competing interest.

